# Empirical determinants of adaptive mutations in yeast experimental evolution

**DOI:** 10.1101/014068

**Authors:** Celia Payen, Anna B. Sunshine, Giang T. Ong, Jamie L. Pogachar, Wei Zhao, Maitreya J. Dunham

**Author notes:** Corresponding author: Maitreya J. Dunham Box 355065 Department of Genome Sciences, University of Washington, Seattle, Washington 98195. Major research organism: *Saccharomyces cerevisiae.

## Abstract

High-throughput sequencing technologies have enabled expansion of the scope of genetic screens to identify mutations that underlie quantitative phenotypes, such as fitness improvements that occur during the course of experimental evolution. This new capability has allowed us to describe the relationship between fitness and genotype at a level never possible before, and ask deeper questions, such as how genome structure, available mutation spectrum, and other factors drive evolution. Here we combined functional genomics and experimental evolution to first map on a genome scale the distribution of potential beneficial mutations available as a first step to an evolving population and then compare these to the mutations actually observed in order to define the constraints acting upon evolution. We first constructed a single-step fitness landscape for the yeast genome by using barcoded gene deletion and overexpression collections, competitive growth in continuous culture, and barcode sequencing. By quantifying the relative fitness effects of thousands of single-gene amplifications or deletions simultaneously we revealed the presence of hundreds of accessible evolutionary paths. To determine the actual mutation spectrum used in evolution, we built a catalog of >1000 mutations selected during experimental evolution. By combining both datasets, we were able to ask how and why evolution is constrained. We identified adaptive mutations in laboratory evolved populations, derived mutational signatures in a variety of conditions and ploidy states, and determined that half of the mutations accumulated positively affect cellular fitness. We also uncovered hundreds of potential beneficial mutations never observed in the mutational spectrum derived from the experimental evolution catalog and found that those adaptive mutations become accessible in the absence of the dominant adaptive solution. This comprehensive functional screen explored the set of potential adaptive mutations on one genetic background, and allows us for the first time at this scale to compare the mutational path with the actual, spontaneously derived spectrum of mutations.

**AUTHOR SUMMARY:** Whole genome sequencing of thousands of cancer genomes has been conducted to characterize variants including point mutations and structural changes, providing a large catalogue of critical polymorphisms associated with tumorigenesis. Despite the high prevalence of mutations in cancer and technological advances in their genotyping, cancer genetics still presents many open questions regarding the prediction of selection and the functional impact of mutations on cellular fitness. Long term experimental evolution using model organisms has allowed the selection for strains bearing recurrent and rare mutations, mimicking the genetic aberrations acquired by tumor cells. Here, we evaluate the functional impact of thousands of single gene losses and amplifications on the cellular fitness of yeast. Our results show that hundreds of beneficial mutations are possible during adaptation but not all of them have been selected in evolution experiments so far performed. Together, our results provide evidence that 50% of the mutations found in experimentally evolved populations are advantageous, and that alternative mutations with improved fitness could be selected in the absence of the main adaptive mutations with higher fitness.

**BLURB:** A combined view of potential adaptive mutations, generated by a systematic screening approach, coupled with the mutational spectrum derived from experimentally evolved yeast reveals the usage of accessible evolutionary solutions.

## INTRODUCTION

Whole genome sequencing of thousands of human tumors has uncovered a huge number of variants including point mutations and structural changes, providing a large catalog of mutated genes across all major cancer types [1-4]. Recent advances in profiling initiatives and systematic genomic analysis of tumors have identified novel mutated genes and rearrangements, raising the prospect of discovering new important drivers of tumorigenesis [2]. However, another recent study discovered that within the list of putatively significant genes, the number of false-positives is also increasing [5]. Given the vast number of mutations identified in most tumors, determining the functional impact of each mutation is a daunting task. The most frequently used approach in cancer genetics to identify the few driver mutations among the many mutations that don’t affect fitness (often called passenger mutations) relies on the hypothesis that genes and pathways important for the development of the disease are recurrently mutated in independent tumors. Those candidate driver genes can then be tested experimentally. Based on such predictions, genes responsible for cell proliferation, drivers of oncogenesis, cell survival, cell cycle, invasion and drug resistance have been identified using RNAi and pools of short hairpins in nematodes and mammalian cell cultures [6,7,8]. While informative, these approaches have not yet been able to assess in an unbiased way the full contribution of mutations to the genetic basis of cancer initiation.

Within the microbial experimental evolution research community, there is a similar need to identify loci contributing to adaptation (also known as adaptive mutations) in the growing list of mutations identified in laboratory-evolved populations. Several recent Evolve and Resequence studies [9], where populations or clones have been sequenced after adaptation to a specific condition, have dramatically increased the list of mutations associated with adaptation in different conditions [10-18]. Within this rapidly increasing dataset, only a few mutations have been fully characterized with regard to function. Similar to studies investigating human disease candidate genes, large-scale studies from the microbial community have distinguished adaptive mutations from background neutral mutations on the basis of statistical approaches such as frequency, enrichment and recurrence [10,11,17,19-23].

Despite sophisticated genetic systems, dissecting the functional consequences of every mutation observed in a population is still tedious, though generally experimentally straightforward. For example, simple genetics can be used to reassort mutations, followed by fitness characterization of segregants carrying individual mutations. This strategy has been performed on a few evolved clones and has demonstrated that evolved clones isolated after several hundred generations of propagation in nutrient-limited chemostats carry 1 to 2 adaptive mutations [24] [Sunshine et al, submitted, See Supplementary file]. *Saccharomyces cerevisiae* is particularly well suited for determining the relationship between genetic variation and fitness at genome scale. Ideally, the functional effects of every possible mutation should be tested. Since recreating and annotating all possible mutations is not yet feasible, the field has instead created systematic dosage series to mimic the most common mutations such as loss- or gain-of-function (LOF and GOF) and deletion or duplication of genes [25-29]. While mimicking LOF, GOF, deletion and duplications, those collections doesn’t take into account mutations that would not be mimicked by copy number changes, such as specific protein coding mutations that generate new activities or more subtle loss of function effects than full knockout alleles. Despite the large number of studies that have used these barcoded collections to detect haploinsufficiency, dosage sensitive genes, synthetic lethality, drug-sensitive mutations, and a huge number of other phenotypes [27,30-38], only a few studies have looked at beneficial mutations (mutations that increase fitness). For example, one study quantified antagonistic pleiotropy in a variety of laboratory conditions and determined that while 32% of deletion strains are less fit than a wildtype reference, only 5.1% of the strains were more fit [39]. Another study identified a large number of heterozygous deletion mutations as being beneficial, but also demonstrated that haploproficiency was context-dependent [26].

Most of these studies have used phenotype data as a way to investigate gene function. However, we can also approach these data from an evolutionary genetics perspective: the ability to identify beneficial mutations *en masse* allows us to survey the set of beneficial mutations upon which adaptive evolution acts. Knowing this landscape allows us to address a number of open questions: what is the distribution of fitness effects of mutations, and how does this distribution compare for loss of function *vs*. gain of function mutations? Which of the possible beneficial mutations are actually utilized by evolution? Are these usage patterns driven strictly by the hierarchy of mutation fitness, or do other factors affect which mutations are observed? How much does the distribution of adaptive mutations differ among different genotypes or selective conditions? For example, how do haploids and diploids differ in the available pool of beneficial mutations, and how might such differences affect the paths by which adaptation can proceed? Finally, to what degree can evolution be perturbed to follow new paths?

The goal of our research was to address these questions using a paired functional genomics and experimental evolution system. We first created a near-comprehensive single-step mutations list by measuring the fitness of almost all single *S. cerevisiae* gene deletions and amplifications. We accomplished this using pooled competition of thousands of mutants in nutrient limited chemostats combined with barcode sequencing. We found that while most single gene copy number changes are neutral or negatively affect fitness, ∼600 mutations increased fitness and correspond to potential evolutionary solutions. We next compared the single-step mutation fitness to the actual mutation spectrum derived from experimental evolution studies performed in this study and also collected from the literature. We found that 50% of the mutations are predicted to positively affect fitness. In sulfate-limited condition, mutations in one gene dominate both the single-step fitness landscape and the observed mutational spectrum, while in the two other conditions the increase in fitness is driven by a large number of beneficial mutations of smaller effect size. Finally, we show that these constraints can be modified by eliminating the highest fitness paths, upon which the evolving cultures explore alternative beneficial mutations.

## RESULTS

### A comprehensive survey of first-step mutations

Pooled competition experiments followed by an Illumina-based barcode sequencing method have been used to accurately determine the fitness effects of hundreds of pooled mutants [25]. We performed a dose-response curve for ∼80% of all the genes of the yeast genome in three different nutrient-limitations using five different yeast barcoded genomic collections (**Table S1**) outlined as follows: two deletion collections in which each gene is replaced by a selectable marker and a unique DNA barcode in one haploid and one heterozygous diploid background [34], one control collection where thousands of unique barcodes have been placed at a single known neutral genomic location [35], and finally two collections of diploid strains bearing plasmids where each gene and its native promoter has been cloned into a barcoded plasmid present at either low or high copy [32,33]. A schematic description of the method is presented in **Figure 1**. While the individual elements of the methodology used here are well established in the literature, this work is the first attempt to compare spontaneously derived mutations from experimental evolution with the potential set of adaptive mutations discovered using a systematic genetic screen.

**Figure 1.**
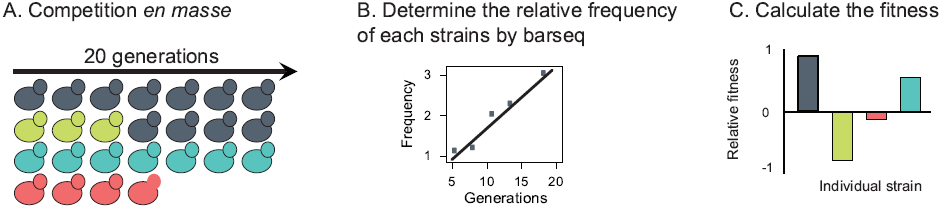
Experimental design for genome-wide pooled competition experiment. The proportion of each strain was measured every 3 to 4 generations during a pooled competition assay, in which all strains from one collection were mixed together in the same ratio and grown at steady state for 20 generations (**A**). The frequency of the corresponding barcode at each time point was measured using the barseq method (**B**), and the fitness of each strain computed (**C**).

Using the five pools described above, we conducted a total of thirty screens in three previously explored chemostat culture conditions (phosphate-limitation, glucose-limitation and sulfate-limitation). The proportion of each strain was measured during a pooled competition assay, in which all strains from one collection were mixed together at the same abundance and grown for ∼20 generations (**Figure S1**). To overcome stochastic effects due to drift, we used cultures of large population size (∼10^9^ cells), as this strategy has been a successful way to maintain diversity [26]. The pooled competitions were performed during a very short period of time (20 generations) to limit the effect of *de novo* mutations occurring during population growth. While other studies have been able to quantify fitness effects of mutations from as few as two time-points, we sampled the mixed population every three generations to maximize the accuracy of the fitness quantification. The frequency of each strain at each time point was measured using barcode sequencing (barseq) (**Figure S2**) [25].

### The functional screening of mutations uncovers hundreds of accessible adaptive mutations

We quantified a total of 100,853 relative fitnesses ranging from -36.5 to +42.8% based on an average of 462 reads per gene per competition and created an experimental fitness landscape of single gene copy number change from four different yeast collections in three conditions (**Figure 2** - **Table S2**). Mutants of 2,133 genes were measured in all twelve experiments (three conditions and four collections), with an additional 2,953 genes sampled by at least one experiment. To determine the inherent noise in our experimental system, which could originate from strain construction, pool generation, competition and/or sequencing, we first quantified the relative fitness of ∼2,000 isogenic barcoded wild-type strains pooled and competed in the same way as for the other four strain collections. As expected, the fitness distributions of these mutants were tightly centered on 0 (**Figure 2** - **Table S3**). We then used the maximum and minimum fitness difference detected in the control pool as conservative cutoffs (±10%) to determine which strains from the four other collections had a strong fitness benefit or deficit when compared to the wild type strains. This cutoff also corresponds nicely with analysis from Otto based on similar evolution experiments performed by Paquin and Adams [40] demonstrating that a beneficial mutation with a 10% fitness increase will reach 5% of the population in ∼200 generations and will fix in ∼500 generations [41]. This analysis suggests that mutations causing less than 10% fitness increase will rarely be observed in our experimental evolution timescale. The functional screening of pooled mutants revealed that most of the mutants display wild-type fitness. Using the 10% cut-off, we detected an enrichment of mutants with a decreased fitness (n=1693 *vs*. 19 for the control pool) and an increased fitness (n=506 *vs*. 80 for the control pool) respectively compared to the control pool (Chi square, *p* <0.0001) (Figure2).

**Figure 2.**
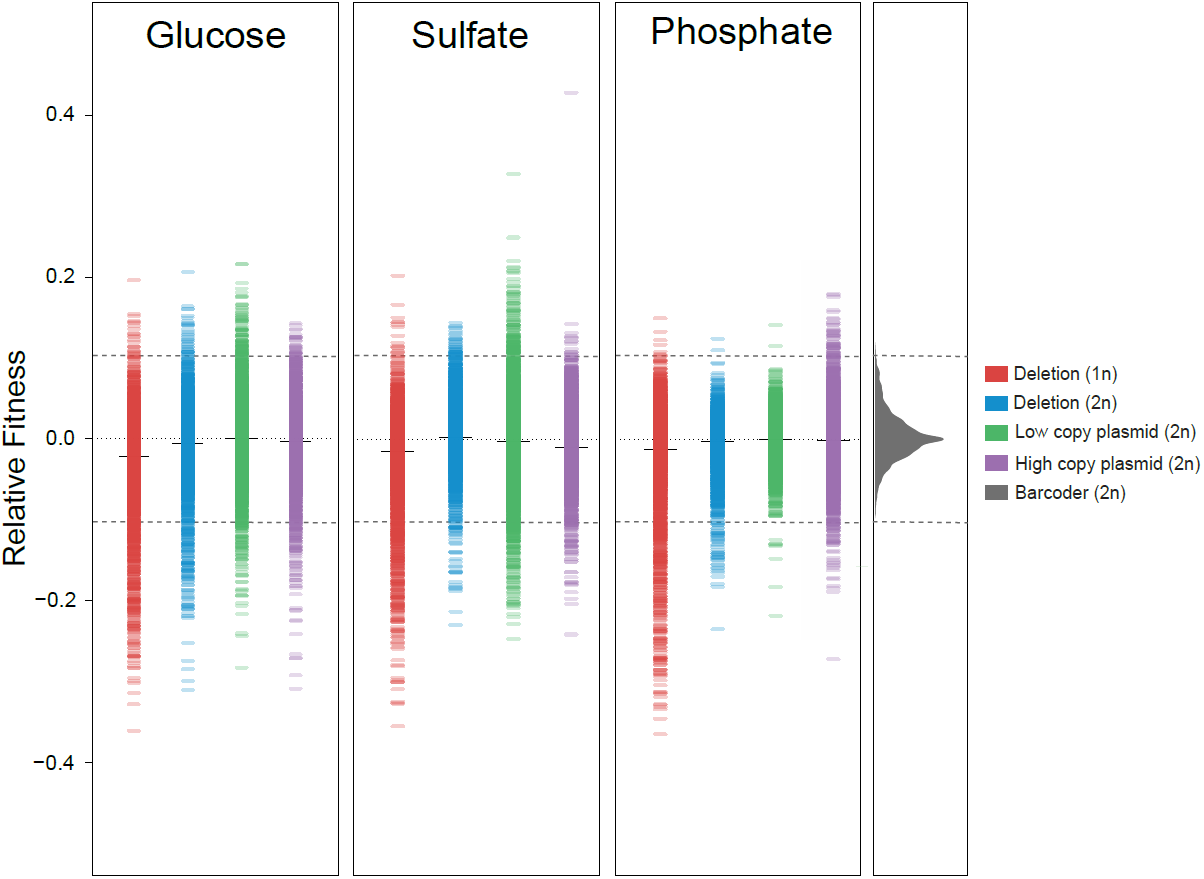
Distribution of the fitness effects of single gene amplification and deletion. Distribution of the fitness measurements of the deletion collections and the plasmid collections in three conditions: glucose-limited, sulfate-limited and phosphate-limited chemostats. The fitness of each strain is shown as small line or as a distribution for the control collection (in grey). The thick black line represents the mean. Dashed grey lines indicate the cut-off of ±10% measured using control pooled collection.

We focused first on the 506 mutants showing increased fitness, hypothesizing that mutations affecting these genes would be more likely to be adaptive during growth under strong selection. Despite making up just 47% of the mutations tested, 73% of the beneficial mutations we detected are from the plasmid collections where the gene copy number is increased, suggesting that in diploids, gain-of-function mutations and duplications are more likely to produce fitness gains than are loss-of function mutations. Among the genes associated with a fitness increase, *SUL1* was notable with the highest fitness measure (42.8% in sulfate-limited condition for a strain carrying a high copy number plasmid). We previously demonstrated that the amplification of this gene is recurrently selected during experimental evolution in sulfate limitation, and that increasing the copy number of *SUL1* via expression on both low and high copy number plasmids results in a fitness improvement [42,43]. Our screen detected a putative secondary adaptive mutation in the vicinity of *SUL1* on chromosome II: *BSD2*, a gene involved in the downregulation of the metal transporter proteins, Smf1 and Smf2 [44,45]. The amplification of *BSD2* increases the fitness of the cells by 5% and 12.4%when amplified in sulfate- and glucose-limitation conditions respectively. In our previous studies of the *SUL1* amplicon, we detected only three independent clones where the *SUL1* amplicon excluded the gene *BSD2.* The fitness of each of the 13 strains harboring an amplification containing both *SUL1* and *BSD2* is higher than the fitness of three strains with an amplicon containing only *SUL1* but not *BSD2* [43]. Reintroducing *BSD2* into one of the three strains using a low copy plasmid increased the fitness by 5% (37.7% to 43.8%), demonstrating that the functional screen with pooled strains is a reliable method to detect small effect and secondary adaptive mutations, and suggesting that the two mutations have an additive effect on the fitness.

Our functional screens revealed the presence of hundreds of possible beneficial mutations (223 in sulfate-, 210 in glucose- and 73 in phosphate-limited conditions). We next sought to apply the functional knowledge gained from the genome-wide analyses described above to the hundreds of *de novo* mutations identified in laboratory evolution experiments. Using this combined dataset, our goal was to ask which particular adaptive mutations are selected and why.

### Mutational spectrum in microbial evolution experiments

To determine the mutational signature of adaptation using laboratory evolution, we sequenced and detected 150 mutations in 16 populations and 34 clones of both haploid and diploid yeast evolved for over 100 generations (122 to 328) under conditions identical to those in which our functional screens were performed (six sulfate-, six phosphate- and four glucose-limited chemostats) [42] (See **Materials and Methods**). To explore this question further, we also collected a large set of mutations from various Evolve and Resequence studies performed under a variety of conditions in yeast [10-12,16,43,46]. Not all the conditions overlap with our functional screens, but they are useful for cross-condition comparison. In total, we compiled 1,167 mutations in 1,088 genes from 106 long-term laboratory evolution experiments conducted in eleven different conditions from nine published studies in addition to this one (**Table S4**). The features of these studies and the resulting mutations are summarized in **Table 1**. The complete list of mutations, their frequencies, and their predicted effects are given in **Table S4**. The compiled mutation catalog does not take into account chromosomal rearrangements, as these events were not always measured in the different studies.

**Table 1.** Mutational catalog subdivided by conditions ploidy and sample

|  |  | Number of mutations | This study<sup>1</sup> |  | Table 1: Mutational catalog subdivided by conditions ploidy and sample |
| --- | --- | --- | --- | --- | --- |
| Conditions | YPD | 720 | NA |  |  |
|  | Glucose | 224 | 23 | 2 |  |
|  | Sulfate | 97 | 76 |  |  |
|  | Phosphate | 54 | 51 | 3 |  |
|  | Other | 72 | NA | 4 |  |
| Ploidy | Haploid | 1017 | 75 |  |  |
|  | diploid | 150 | 75 |  |  |
| Sample | Clones | 305 | 75 |  |  |
|  | Population | 862 | 75 |  |  |

### LOF mutations are enriched in haploids and are depleted and recessive in diploids

Comparing the mutational spectrum across many environments, strains and ploidies allows us to extract mutational signatures and infer the properties of beneficial mutations in yeast. Ploidy in particular has been a subject of much interest since the observation that haploids and diploids adapt at different rates [40]. Two recent studies have shown that loss-of function mutations were commonly selected in evolved populations of haploid yeast [10,11]. Based on a small number of mutations tested in diploids, another study concluded that mutations affecting cis-regulating regions are co-dominant in heterozygous diploids [47]. Though these results are suggestive, because no other Evolve and Resequence studies have been performed in a diploid background, too few data have previously been available to draw firm conclusions about how the mutational landscape differs by ploidy.

We divided these mutations into four groups based on SNPeff, an annotation program that predicts the functional impact of the mutation of a gene, as follows [48]: (1) high impact mutations such as frameshifts and the gain or loss of a start or stop codon; (2) moderate impact such as non-synonymous site changes and the deletion or insertion of a codon; (3) low impact synonymous mutations; and (4) modifiers, corresponding to mutations 5’ of a gene, in intergenic regions and in introns. We found that the mutation signature is different between haploid and diploid strains (Fisher exact test, p<10^-16^, corrected for multiple testing). In haploids the main category of beneficial mutations is LOF by gain of a stop codon (Chi square, *p*=0.003, Table 2) consistent with a previous finding that LOF mutations dominate in experimental evolution of haploid yeast [11]. In contrast, LOF are depleted in diploid strains, which instead show an enrichment for intergenic and 5’ upstream mutations, suggesting that amplification and GOF mutations may be more important in this background (Chi square, *p*<10^-4^, Table 2). This result is consistent with our previous observations that evolved diploid strains contain more and larger gene and chromosome copy number variants than evolved haploids. We next investigated if the difference between haploid and diploid was a general rule across environments. Using only mutations discovered in haploid and diploid strains evolved under matched conditions, we detected that the mutational signature was different between haploids and diploids in glucoselimitation (Fisher exact test, *p*<10^-14^) with an enrichment of LOF in haploid (Chi-square, n= 224, *p*<10^-9^), but only a slight tendency is observed in phosphate-limitation (Fisher exact test, n=54 *p*=0.053) and none in sulfate-limited conditions (Fisher exact test, n=100, *p*=0.72). The difference between ploidies is likely explained by the tendency of LOF mutations to be recessive [49] compared to mutations that increase gene expression, which may be more likely to have an effect as a heterozygote. Though loss-of-heterozygosity has been observed in diploid populations [42,49], these are relatively rare. To test this directly, we determined how many LOF mutations detected as beneficial in a haploid context might lose this effect when heterozygous in a diploid. We compared 58 beneficial mutants from the haploid deletion collection to the fitness of the heterozygous diploid mutants and found that these mutations do show a tendency to be recessive, with the average loss of fitness between haploid and diploid of 8.6%. Only nine genes showed no statistical change in fitness, indicating that a subset of LOF mutations can in fact be dominant (*WSC3*, *TIM12*, *IPT1*, *MMS22*, *UFO1*, *NDL1*, *PBS2*, *YGR051C* and *YLR280C*). We also found that the distribution of mutations is not uniform along the coding sequence. Disruptive mutations (high impact mutations) are enriched near the start codon (Wilcoxon rank-sum test, p<10^-3^ **Figure S3A**), indicating a preference for early truncations, which are most likely to cause severe LOF.

**Table 2.** Comparison of the mutational signature in haploid and diploid strains

| Class | All | diploid | haploid | pvalue* | qvalues | SNPeff |
| --- | --- | --- | --- | --- | --- | --- |
| Stop gained | 123 | 5 | **118** | 0.003 | 0.005 | High |
| Start lost | 8 | 1 | 7 | 0.97 | 0.591 | High |
| Stop lost | 2 | 0 | 2 | 0.59 | 0.469 | High |
| Frameshift | 6 | 0 | 6 | 0.74 | 0.496 | High |
| Non-synonymous | 817 | 97 | 720 | 0.15 | 0.201 | Moderate |
| Codon deletion/insertion | 3 | 0 | 3 | 0.51 | 0.469 | Moderate |
| Synonymous | 142 | 15 | 127 | 0.46 | 0.469 | Low |
| 5'upstream | 13 | **7** | 6 | 0.0001 | 0.002 | Modifier |
| Intron | 5 | 1 | 4 | 0.63 | 0.469 | Modifier |
| Intergenic | 48 | **24** | 24 | 0.0001 | 0.002 | Modifier |
| Sum | 1167 | 150 | 1017 | 2.5e-315 | 1.68e-314 |  |

\* Chi-square and Fisher's exact test *p*values

### Mutational pathways are constrained

Recurrence-based models, which assume oncogenes are recurrently mutated in several tumor samples, are still one of the most widely used approaches to identify putative driver genes in cancer [50-52]. The repeatability of adaptive trajectories has also been extensively observed in the microbial research community and has led to the discovery of drivers of adaptation such as *SUL1*, *HXT6/7*, and *RIM15* in *S. cerevisiae* and *rpoS* in *Escherichia coli* [11,19,20,42,53]. Of the 1,088 genes mutated in the catalog we compiled, 154 genes were found with a mutation in more than one sample, and among them 19 genes were found mutated more than five times independently (**Figure 3A**). We detected that recurrently mutated genes are highly enriched in mutations categorized as high impact (Fisher exact test, *p*<10^-16^) (**Figure 3B**) and are longer than genes with only one hit (Wilcoxon rank-sum test, *p*<10^-16^) (**Figure S3B**). In order to detect true adaptive mutations and discard false-positives, several studies have developed tools to correct for gene length [5]; in our study we decided instead to attempt to infer the functional impact of mutations on cellular fitness using our screen results.

**Figure 3.**
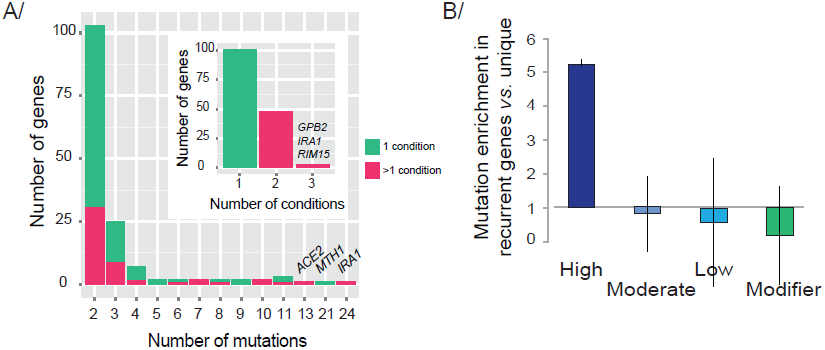
Recurrently mutated genes reveal how evolution is constrained. **A-** Repeatability of adaptation and parallelism at the gene level. Genes classified by number of mutations detected during Evolve and Resequence studies. 154 genes were found to be hit by more than one mutation. 48 recurrent genes were found mutated in more than one conditions (small panel). **B-** Enrichment of high impact mutations in recurrently mutated genes when compared to genes found with only one mutation. Error bars are 95% CI.

### Prediction of evolutionary response to strong selection

Despite the presence of more than one hundred recurrent mutations, a large number of genes are mutated in only single populations. Since the number of Evolve and Resequence experiments is currently still relatively small, akin to a non-saturating genetic screen, adaptive mutations are likely to be found in the class of singletons and would be missed by a recurrence method. As an alternative strategy toward specifically identifying adaptive mutations, we compared the mutations found in the evolution experiments with known beneficial mutations identified by our genomic screen.

From the functional screen described above, we detected 506 beneficial mutations targeting 458 genes; among them, 86 genes were found with a hit in our compiled mutation catalog, 27 in their corresponding conditions and the rest in the other conditions (YPD and nitrogen-limited). We also detected 21 recurrently mutated genes present in the list of beneficial mutations (**Table S5**). From the mutational catalog, 41 of 70 recurrent mutations were not associated with beneficial fitness in matched conditions in our functional screen. A third of them are not present in the mutant collections; another third were selected during experimental evolution performed in more than one condition and might not represent true convergent adaptation; and eight of them have a fitness ranging from 3 to 9%, below our stringent threshold for significance (*PDE2*, *LCB3*, *SSK1*, *DAL81*, *RAS2*, *MTH1*, *IRA1* and *RGT1*). The remaining five genes were recurrently mutated, but had no obvious benefit in their given conditions (*VPS25*, *MNN4*, *FRE5* and *GSH1* in glucose and *PHO84* in phosphate). One example, *MNN4* has been found mutated in two independent populations grown in glucose limitation; however we measured no fitness benefit in our functional screen and no fitness benefit was reported in a competitive assay using evolved clones [24]. These five genes could be recurrently mutated by chance, or fitness increases caused by these mutations are not mimicked by gene amplification or deletion collections, which may be the case for partial loss of function mutations or gain of function mutations that create a new activity. Alternatively, these mutations may only have a benefit in a specific genetic background. This data show that convergent evolution cannot be used as the only parameter to predict evolutionary outcomes and more comprehensive and unbiased detection of adaptive mutations requires a more direct method such as functional screening.

### 50% of the mutations accumulated during experimental evolution are adaptive

Next we wanted to determine how many adaptive mutations were carried by each sequenced population and clone, using the frequency of recurrence combined with data from the functional screen. We determined that 91% of the samples (clones and populations) carried at least one predicted driver mutation. Of these samples, each contained an average of 5.2 confirmed known beneficial mutations: 7.6 per population and 2 per clone, with an average of 0.47 adaptive mutations per total mutation (**Figure 4A** - **Table S6**). No difference was detected between conditions (**Figure 4B** - **Table S6**). Three populations with no predicted beneficial mutations were cultivated in nitrogen-limiting conditions. However, these strains have been shown to carry Copy Number Variants (CNVs) [12], and we did not include nitrogen limitations in our functional screen. We also detected 24 mutations from the experimental evolution studies in genes that are associated with deleterious mutations based on our functional screen performed in the same conditions. However, none of the mutations were predicted to have a high impact on the function of the gene, and so they might instead be neutral or near-neutral passenger mutations. Thus, combining functional screening of mutations and whole genome sequencing of populations and clones in this way, we are able to identify both drivers of adaptation and also unexplored fitness peaks. We conclude that evolution is partly predictable based on the repeatability of adaptive trajectories in independent evolution experiments and reflects at least in part the underlying fitness distribution of possible mutations.

**Figure 4.**
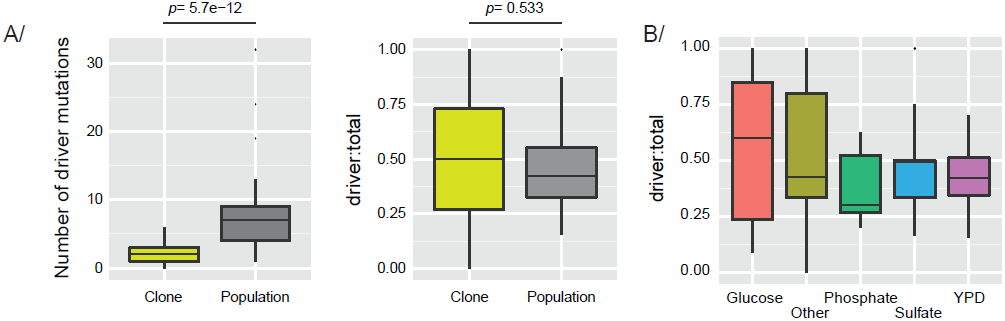
Driver mutations. **A-** Boxplot representing the number of driver mutations and the ratio of driver to total mutations detected in evolved clones and populations. The significance of the difference between clones and populations was estimated using Wilcoxon-ranked test. **B-** The ratio of driver mutations to mutation total is not conditions specific (*p*=0.28; 0.70; 0.36; 0.78 and 0.36 for glucose-limited; sulfate-limited; YPD; other and phosphate-limited respectively).

### The set of beneficial mutations reveals potential drivers of adaptation

The analysis above defines the subset of adaptive mutations actually utilized by experimental evolution. However, the screen for beneficial mutations identified a large mutational reservoir with many additional accessible evolutionary paths [54]. To determine what differentiates the actual mutation spectrum from the potential mutation pool, we excluded the mutations that had already been identified in experimental evolution, and found 369 potential adaptive mutations that were unobserved in the existing evolved populations. Given the population size of the cultures used for experimental evolution (10^5^ to 10^10^ cells depending on the experimental set-up), the number of generations grown (50 to 1000 generations), and the size of the yeast genome (∼12 megabases), every base mutation must have been explored many times in the ensemble of experiments.

We used our functional screen to determine whether the mutations actually selected for during experimental evolution differed from the potential adaptive mutations that were never recovered. We detected a statistical difference between the fitness of the beneficial mutations observed in *versus* absent from the experimental evolution studies in glucose-limitation (Wilcoxon rank-sum test, *p*=0.02) but not in phosphate-limitations (Wilcoxon rank-sum test, p=0.6) or sulfate-limitations, which are dominated by the fitness increase caused by *SUL1* amplification (Wilcoxon rank-sum test, *p*= 0.06 and *p*=0.34 in the absence of *SUL1*) (**Figure 5A**). The small number of mutations detected in populations evolved under sulfate and phosphate-limitations (n=94 and 54) may have limited our ability to detect a similar fitness differential as observed in glucose-limitation (n=224). This would suggest that the observed mutation spectrum is driven by the fitness of potential beneficial mutations. The observed mutation spectrum could also be biased away from the highest fitness mutations by differences in mutation rate, as previously proposed [10]. Likely, the lack of mutations in these genes may be the result of a combination of all of these factors, including random chance, epistatic interactions between mutations, and/or a reflection that the pool experiment does not adequately recapitulate the fitness of the *de novo* mutations. Clonal interference is also likely to play a large role. Consistent with previous findings, *SUL1* dominates in the functional screen and in the mutational spectrum (**Figure 5B**), but other highly beneficial mutations (>20% fitness increase) such as mutations in *MAC1* and *PHO3*, two genes coding proteins implicated in copper and phosphate-sulfate metabolism, respectively, are also potential drivers but are never recovered (**Figure 5B**) [10,55]. Conversely, in glucose limitation, many beneficial mutations of similar fitness are possible, and so more variety in outcomes and broader sampling of the mutational reservoir is observed.

**Figure 5.**
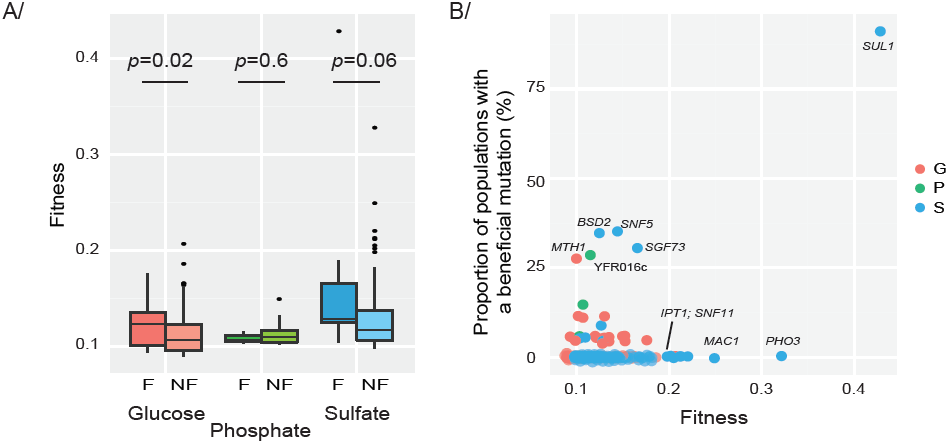
Alternative accessible evolutionary paths. **A-** The fitness of beneficial mutations found (F) in Evolve and Resequence studies is statistically significantly higher than the fitness of beneficial mutations not found(NF) in glucose-limitation but not in phosphate-limitation and sulfate-limitation. The significance of the difference between the two boxplots for each condition was estimated usinga Wilcoxon-ranked test. **B-** Each point represents the fitness of a strain and the proportion of Evolve and Resequence samples with the corresponding gene mutated. *SUL1* dominates the fitness and mutational spectrum. Several mutations have a high fitness but have never been detected in Evolve and Resequence studies and might correspond to potentialdrivers of adaptation.

### Mutational spectrum in the absence of the main adaptive mutation

To investigate the discrepancy we observed between the single-step fitness landscape and the observed mutational spectrum, and to test the predictability of experimental evolution, we wanted to test if we could detect unobserved adaptive mutations by inhibiting the selection of the main driver of adaptation. We have shown in previous work that *SUL1* amplification dominates the mutational spectrum [42,43] and is the mutation with the highest fitness in our screen (**Figure 5B**). Additional adaptive mutations might be undetectable in sulfate-limited conditions due to the presence of such a strong fitness peak. We hypothesized that by eliminating the selection of the *SUL1* amplification, a variety of smaller effect mutations will be selected, an outcome more similar to the pattern observed in glucose-limitation. To explore the mutational landscape in the absence of the main adaptive mutation, we screened two evolved populations in which no *SUL1* amplification was detected by qPCR (**Figure 6A**) and aCGH (data not shown) even after 200 generations of cultivation in sulfate-limited conditions. The fitness of the clones and populations without *SUL1* amplification (∼30%) (**Figure 6B**) are on the lower end of the fitness range of previously studied evolved clones with *SUL1* locus amplifications (37% to 53%) [43]. To establish which mutations were responsible for this phenotype, we performed whole genome sequencing and called SNPs and INDELs of the clones and the populations isolated at generation 200. One nonsense mutation was detected in the previously identified adaptive gene *SGF73* for one of the clones (**Table S4**). Two independent non-synonymous mutations (N263H and N250K) in the coding-region of *SUL1* were also detected in both populations. Wild type strains containing those mutations were created and we detected a fitness increase of 23.1% (±2.3) for the strains carrying N250K and 17.7% (±1.22) for the strain carrying N263H. In addition, for the second clone, we detected a 5.1 kb deletion on chromosome IV (4.8kb, 587839592999) affecting four genes (*FMP16*, *PAA1*, *IPT1* and *SNF11*) (Figure 6C). From our functional screens, we found that deletions of *IPT1* and *SNF11* are beneficial in glucose and sulfate-limited conditions (10 to 20% fitness increase) but mutations in these genes were never detected in any of our previous evolved populations (**Figure 5B**). Since these genes are adjacent on the chromosome, we suspected that one of these genes may be a false positive, resulting from a known artifact called the neighboring gene effect [56]. To decipher which deletion drives the increased fitness, we used complementation screens using centromeric plasmids, and found that the deletion of either gene drives the fitness increase in the evolved strain (**Figure 6D**). *SNF11* is a subunit of the SWI/SNF chromatin remodeling complex, which is known to act as a tumor suppressor in humans [57], while *IPT1* is implicated in the metabolism of membrane phospholipids and nutrient intake [58]. Thus, small-effect mutations detected in the functional screen are relevant although they may not be detected at first in experimental evolution. We predict that additional evolution experiments that remove the *SUL1* amplification path would eventually explore even more alternative accessible evolutionary routes.

**Figure 6.**
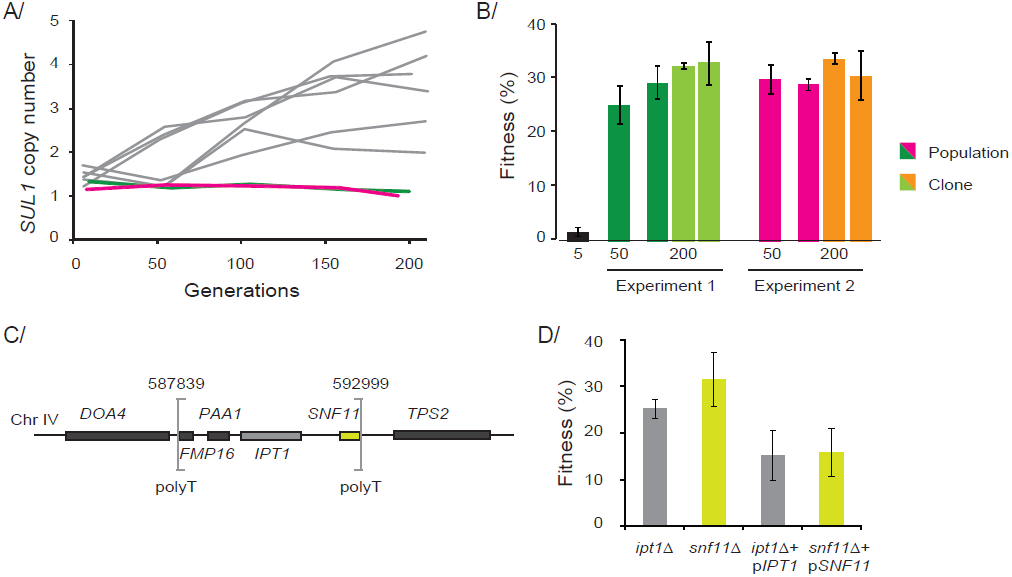
New beneficial mutations are selected in absence of the main driver. **A-** The copy number of *SUL1* was assessed using qPCR analysis on samples taken from two independent experiments in which *SUL1* did not amplify (green and pink), compared with previously published data from wild type strains (in grey) (Payen et al. 2014). **B-** Fitness coefficient of population samples at generation 5, 50 and 200 and the fitness of two clones isolated at generation 200. **C-** Small deletion (∼5kb) detected in population from experiment 2 on chromosome IV encompassing four genes (between brackets), polyT sequences are present at the breakpoints, the color of the boxes represent the orientation of the genes (yellow: gene on the Watson strand, grey: genes on the Crick strand). **D-** Fitness coefficient of the two deletion strains *lpt1*Δ and *snf11*Δ A, and both deletion strains complemented with *IPT1* or *SNF11* on a low copy plasmid in sulfate limited condition.

## DISCUSSION

Our work addresses a central topic in evolutionary biology, the relationship between genotype and fitness and how evolution is constrained despite the presence of alternative accessible paths. For this purpose we generated both a nearly complete set of possible beneficial mutations and a catalog of mutations actually observed during long term experimental evolution. Using those two datasets, we were able for the first time to compare potential and actual beneficial mutations and begin to understand why some mutations are selected or not.

### Patterns and reproducibility of evolution

By compiling a catalog of >1,000 mutations identified in 109 independent evolution experiments from this study and others (**Table S4**), we were able to ask a variety of questions about the reproducibility of adaptation, and the features of beneficial mutations over multiple conditions and ploidy states. We detected an excess of loss of function mutations in haploids, as previously shown by Kvitek and Sherlock [11]. Moreover we estimated that mutations predicted to modify gene expression level are statistically enriched in diploid compared to the number in haploids. Mutation rate has been shown to be similar in diploids and in haploids [59,60], suggesting differential selection or a mechanism based on genetic context and not on the mutation rate. Several studies have also shown that mutations have a greater effect on the fitness of haploids than heterozygous diploids [61], which we were also able to show, and that the frequency of fixation is higher in diploids [40]. Mutations affecting cis-regulating regions have often been described as co-dominant while most coding region mutations will be recessive [47]. Large CNVs have also been seen to be enriched in diploid backgrounds *versus* haploids [42], suggesting that a diploid context for aneuploidy and CNVs might buffer the detrimental cellular effect seen in haploids [62,63].

In agreement with previous reports [11,17,19,42,53], we detected that selection of mutations under laboratory controlled conditions results in a non-uniformity of the distribution of mutations across the genome, as we detected over one hundred recurrently mutated genes (**Figure 3A**). We detected that the same beneficial phenotype can arise through identical genomic changes (recurrently mutated genes) [10,17] and also through different, apparently unrelated mechanisms as 85% of the genes were only hit once by a mutation. As the recurrence based method offers an unsatisfactory prediction of the impact of mutations on cellular fitness [64], functional screening of all mutations was still required to discriminate neutral and passengers mutations from causative mutations.

### Experimentally surveyed set of beneficial mutations

To solve this problem, we built a nearly complete set of beneficial mutations based on both gain and loss of function of nearly every gene in the yeast genome. This data set was generated by competing libraries of systematically created mutant strains *en masse* and then by analyzing the results by barcode sequencing. The functional screen revealed that most single gene deletions or amplifications did not affect the fitness of the cells, demonstrating the robustness of cellular fitness to subtle genomic changes (**Figure 2**). We also detected 506 mutants with a fitness increase. A large proportion of the beneficial mutations originated from the overexpression collection, revealing that gain-of function mutations positively affect cellular fitness in this background. These data illustrate the large number of accessible adaptive mutations, and allow us to compare this list of potential beneficial mutation with mutations selected during the course of laboratory evolution experiment, in order to ask which mutations are selected and why.

### Evolution is constrained by the fitness of adaptive mutation

By combining the beneficial mutations detected in the functional screening and the mutational spectrum of evolved clones and populations, we were able to determine that 50% of the mutations detected in evolving populations are beneficial. As would be expected, this number is higher than previous estimates of the null distribution of mutation fitness using mutation accumulation lines performed in yeast (6% to 13% of all mutations) [65]. We also found that some mutations dominate the mutational spectrum by dominating the fitness of beneficial mutation. For instance, a particular large effect mutation is nearly always observed in sulfate-limited conditions, while a diversity of smaller-effect beneficial mutations was detected in both glucose and phosphate-limitations.

The comparison also revealed a large number of potential beneficial mutations that have never been observed in any Evolve and Resequence studies so far (**Figure 5B**). We wanted to see if those mutations corresponded to inaccessible evolutionary paths or if they could be selected in some specific conditions. We decided to focus on sulfate-limitation, as one primary evolutionary path is utilized in this condition *(SUL1* amplification). We looked in evolved strains without this mutation, and found that alternative routes could then be explored. The fitness of the evolved population linked to the deletion of two adjacent genes (*IPT1* and *SNF11*).

### Remaining open questions

While producing for the first time at this scale a single-step fitness landscape of single gene mutations in the yeast genome, the functional screen using both amplification and deletion collections has several limitations. The collections available in yeast are based on single gene copy number changes and do not allow study of single point mutations, and protein-coding mutations that are not mimicked by dosage changes, non-genic functional elements or combinations of mutations. To explore the importance of non-genic regions and small genes not present in the yeast collections, billions of individual and combined mutations need to be generated in a comprehensive way, similar to deep mutational scanning of proteins [66], the Million mutation project [67] or by using newly created collection such as the tRNA deletions collection [68] or large telomeric amplicons [Sunshine, submitted, see Supplementary file]. A major challenge now is to identify the combination of genetic variants that modulate the activity of specific pathways. Previous studies in simpler microbial and viral systems have provided evidence for both antagonistic and synergistic epistasis between beneficial mutations [39,69-72]. Synthetic genetic arrays and other similar approaches using the *S. cerevisiae* deletion collection have been used to characterize negative and positive epistatic relationships, and a nearly complete yeast genetic interaction network has been generated using double mutants grown under a single lab condition, showing that genes within the same pathway show similar interaction patterns [73,74]. Further studies with these resources would also allow us to move beyond single gene effects and begin to understand how multiple genes in CNVs and combinations of mutations shape the fitness landscape. By expanding and developing these techniques, the increase of studies combining long term experimental evolution and whole genome sequencing will likely reveal additional subtle mutational signatures and support the causal link between mutations and phenotypes such as the impact of synonymous mutations on gene splicing as has been recently shown in oncogenes [75], and the impact of mutations on cis-regulation in the genome [47,76].

## Conclusions

Our analysis makes clear that the identification of adaptive mutations requires accurate functional screening integrated with variant discovery to allow the confirmation of frequently observed mutations but also the discovery of alternative adaptive mutations. Our results predict that the increase of evolved population sequencing data combined with unbiased and comprehensive functional information to broadly query the genome on a large variety of conditions and genetic backgrounds will result in a more complete characterization of the mutational landscape of adaptation.

## METHODS

**Strains and media used in this study**. The MoBY-ORF collection in *Escherichia coli* was obtained from Open Biosystems and stored at -80C as individual strains in 96-well plates. The plates were thawed and robotically replicated onto LB-Lennox (Tryptone 10g, Yeast extract 5g, NaCl 5g) agar plates containing 5μg/ml of tetracycline, 12.5μg/ml of chloramphenicol and 100μg/ml of kanamycin and grown at 37°C for 14 hours. Colonies were harvested by addition of 5ml LB-Lennox to each plate and subsequently pooled. 50% Glycerol was added and aliquots of 1ml, containing 2x10^9^ cells/ml, were frozen at -80C. Plasmid DNA was prepared from the *E. coli* pool and then transformed into *S. cerevisiae* S288C derivative strain DBY10150 (ura3-52/ura3-52) using a protocol adapted from Gietz and Woods (2005). The yeast transformants were selected on–URA and 200μg/ml G418 plates. 88,756 transformants were pooled together, giving an average library coverage of ∼20x. The MOBY-ORF v2.0 collection was obtained from the Boone lab and crossed for 3 hours with YMD1797 (*MATα*, *leu2∆1*). Clones were selected on MSG/B and G418 (200μg/ml) twice and pooled together. The *MATa/MATα* Magic Marker collection was obtained already pooled from the Spencer lab. The *MATa* Magic Marker library was obtained frozen from the Caudy lab; the strains were selected on -LYS and -MET and pooled together. The barcoder collection was obtained frozen from the Nislow lab. The plates were thawed at room temperature, replicated onto YPD and G418 (200μg/ml) and crossed with FY5 (*MATα*, prototrophic strain), the strains were then selected on MSG/B+G418 (200μg/ml) twice and pooled together. A list of strains used in this study can be found in Table S1.

**Continuous culture in chemostats and pool competition experiments**. Nutrient limited media (sulfate-limited, glucose-limited and phosphate-limited) as described in [19,42,77] were complemented with uracil and histidine (20mg/L) for the Magic Marker pools. The 200ml chemostat vessels were inoculated with 1ml of each pool (∼2x10^7^ cells). Cultures were grown at a dilution rate of 0.17±0.01 volumes/hour at 30°C. We grew the five pools in chemostats for 30 hours in batch and then switched to continuous culture. The cultures reached steady state after ∼10 generations and were maintained for 20 generations in the three conditions (**Figure S1**). A sample taken just after we turned the pump on, was designated Generation 0 (G0), then samples were harvested every 3 generations on average. Samples for cell count and DNA extraction were passively collected twice daily. Each pooled competition was performed in duplicate.

**Genomic DNA preparation, Plasmid extraction, qPCR**. Genomic DNA was extracted from dry, frozen cell pellets using the Smash-and-Grab method [78]. Plasmids from the MoBY collections were extracted with a Qiagen miniprep protocol (QIAprep Spin mini prep kit kit; Qiagen, Hilden, Germany) using the following modification: 0.350mg of glass beads were added to a cell pellet with 250μl of buffer P1 and vortexed for 5min. Then 250μl of buffer P2 was added to the mix of cells and beads and 350μl of buffer N3 was added to the solution, before centrifuging for 10 min. The supernatant was then applied to the Qiagen column following the recommendation of the Qiagen miniprep kit. Plasmid DNA is then eluted in 501μ of sterile water. Smash-and-Grab Genomic DNA was extracted from dry pellet of cells using Smash-and-Grab method and used for barcode verification of single strains using PCR amplification and Sanger sequencing as previously described [43]. For each sample, the plasmid copy number was determined using the copy number of *KanMX* relative to the copy number of *DNF2*, a gene located on chromosome 4 and absent from the two MoBY collections (see **Figure S5**). The primers used are included in **Table S8**. Microarray, whole genome sequencing, SNP calling and qPCR analysis were performed as previously described [43]. Microarray data from this article have been deposited in the Gene expression Omnibus repository under accession GSE58497 (https://www.ncbi.nlm.nih.gov/geo/query/acc.cgi?token=sjgtsgwmdhajdud&acc=GSE58497). The fastq file for each library is available from NCBI Short Read Archive with the accession number PRJNA248591 and BioProject accession PRJNA249086.

**Barseq experiments and fitness measurement.** Amplifications of the barcodes were performed using a modified protocol [25]. Uptag barcodes were amplified using primers containing the sequence of the common barcode primers (bold), a 6-mer tag for Illumina multiplexing (in italics) and the sequence required for attachment to the Illumina flowcell (underlined) (**Table S8**). PCR amplifications were performed in 100μl volume, using Roche FastStart DNA polymerase with the following conditions; 94°C/3min, 25 cycles of 94°C/30sec, 55°C/30sec, 72°C/30sec, followed by 72°C/3min. PCR products were then purified using the Qiagen MinElute PCR Purification kit (cat. No. 28004), quantified using a Qubit fluorometer and then adjusted to a concentration of 10μg/ml. Equal volumes of normalized DNA were then pooled and gel purified from 6% polyacrylamide TBE gels (Invitrogen) using a soak and crush method followed by purification and concentration using Qiagen Qiaquick PCR purification. After quantification using a Qubit fluorimeter, libraries were sequenced using the standard Illumina protocol as multiplexed single read 36-base cycles on several lanes on an Illumina Genome Analyser IIx (GAII). We sequenced thirty multiplexed libraries (UPTAGS only) on several lanes of an Illumina GAII and we obtained on average 25,664,072 million reads that perfectly matched the molecular barcodes per library (**Table S9**). The fastq file for each library is available from the NCBI Short Read Archive with the accession number PRJNA248591 and BioProject accession PRJNA249086 and are listed in **Table S10**. The 6-mer multiplexing tags were reassigned to a particular sample using a custom Perl script (**Supplementary File 1**). Then, each barcode was reassigned to a gene using a standard binary search program (program in C, **Supplementary File 2**). Only reads that matched perfectly to the reannotated yeast deletion collection [25] or MoBY-ORF collection [32] were used. For the barcoder collection, the barcodes were recovered using a compiled list of all barcodes previously published. We were able to recover 1885 barcodes, where1624 barcodes were recovered from the barcode list of the yeast deletion collection and 260 barcodes from the Yeast Barcoders collection [31,35]. Multiple genes with the same barcodes were discarded. The strains with less than 20 counts across the different samples were discarded. The numbers of strains identified for the five collections in the three conditions are summarized in **Table S9**. To avoid division by 0 errors, we added 10 to each barcode count before normalizing to the total number of reads for each sample. To quantify the relative fitness of each strain during growth in the various conditions, we restricted our analysis to when the samples reached ‘steady-state’ phase defined as generations 6 through 20, and used generation 0 as t_0_. The linear regression of the log_2_ ratios between generation 6 and 20 to generation 0 was used to calculate the fitness of each strain and the two replicates measurements were then averaged. The source code is provided in the supplementary materials (R script, **Supplementary File 3**).

**Validation of the fitness measurements and pairwise competition.** To ensure that the pooled fitness measurements accurately reflect the fitness of each strain, we measured the relative fitness of 51 strains from the deletion and plasmid collections that were detrimental, neutral or beneficial, by individual competition against a control strain marked with a fluorescent protein (e*GFP*) in the three conditions used in the pooled experiment. Fitness measurements of individual clones were performed as previously described [43] using FY strains where the *HO* locus had been replaced with *eGFP*(*MATa*: YMD1214 and *MATa/MATα* YMD2196) (**Figure S4**, **Table S7**). The fitnesses are similar in both assays and we observed a strong positive correlation (R^2^=0.83) between the large pool screen and the individual fitness measurements (**Figure S4** and **Table S7**). A second concern is that use of the yeast collections to determine the association of fitness changes could be compromised by mutations or copy number changes preexisting elsewhere in the genomes of the pooled strains. To limit this known artifact, most of the barcoded pools used for these experiments were created either by fresh transformation (in the case of plasmid collections) or from a fresh cross of the commercially available collection stocks with a wild-type strain to dilute any possible passenger mutations (See above in **Materials and Methods**). To avoid de novo mutations achieving high frequency and skewing our fitness measurements, we limited our pooled and pairwise competition to 20-25 generations.

To determine the number of mutations of our validation panel, we screened these fifty-one clones for what is known to be the most common secondary mutation detected in the deletion collection, mutations in the gene *WHI2*, which is involved in the regulation of cell proliferation [25,74]. We confirmed the lack of mutations in the *WHI2* gene in the individual strains by Sanger sequencing (**Table S7**). We also detected no copy number changes at the population level using microarray analysis of the last sample of the competition of the low copy plasmid collection, though this approach would only detect CNVs that achieved at least ∼10% population frequency (data not shown).

## DATA ACCESS

All sequencing data from this study have been submitted to the NCBI Sequence Read Archive (SRA; https://www.ncbi.nlm.nih.gov/sra) under accession number PRJNA248591 and BioProject accession PRJNA249086. Microarray data from this article have been deposited in the Gene expression Omnibus repository under accession GSE58497 (https://www.ncbi.nlm.nih.gov/geo/query/acc.cgi?token=sjgtsgwmdhajdud&acc=GSE58497).

## ACKNOWLEDGMENT

We thank the members of the Dunham lab, members of the Brewer/Raghuraman lab, Matt Rich, Colin McNally, and Joseph Schacherer for helpful discussions and comments on the manuscript. Thanks to Shane Trask for his help with the SRA submission. We are thankful to all of the yeast community who has shared with us several yeast collections, in particular the Boone, Spencer, Nislow and Caudy labs. We thank Can Alkan for assistance with C programs, Loic Paillotin for help with Perl, Ron Hause with ggplot2 and also Charlie Lee from the Shendure lab for assistance with the DNA sequencing. Thanks to Gavin Sherlock and Dan Kvitek for sharing prepublication data.

## AUTHOR CONTRIBUTIONS

Conception and Design: CP, ABS, and MJD; Acquisition of data: CP, GTO, ABS and JLP; Analysis and interpretation of data: CP, ABS, WZ and MJD; Drafting the article: CP and MJD.

## SUPPLEMENTARY FILES

**Figure S1.**
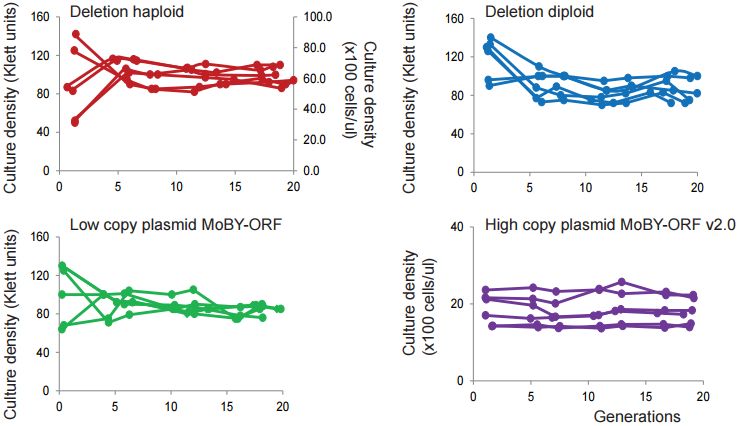
Steady-state in continuous cultures is reached at generation 6. Cell density over time for each pool grown in the chemostat in glucose-limited, sulfate-limited and phosphate-limited for 20 generations.

**Figure S2.**
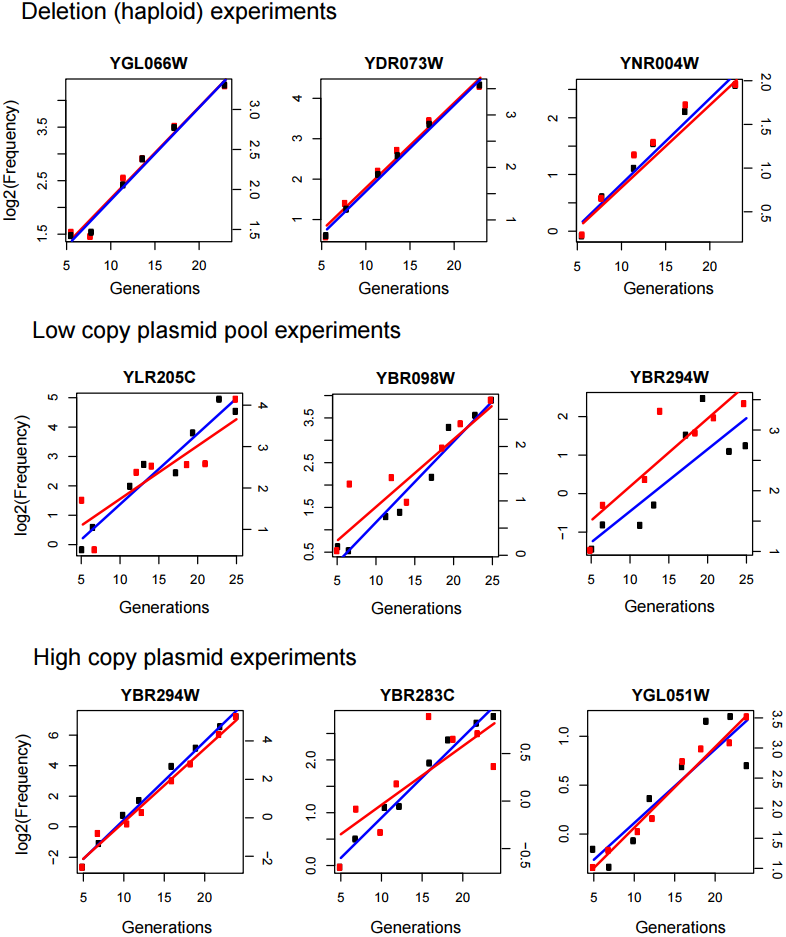
Relative frequency over time of three strains from four collections. Each box, represents the relative frequency of one strain over time, plotted as the log_2_ ratio of the frequency at generation x relative to its frequency at generation = 0 over the ∼20 generations of steady-state competition. Each line, colored blue and red, represents the linear regression used to calculate the relative fitness between generation 6 and 20.

**Figure S3.**
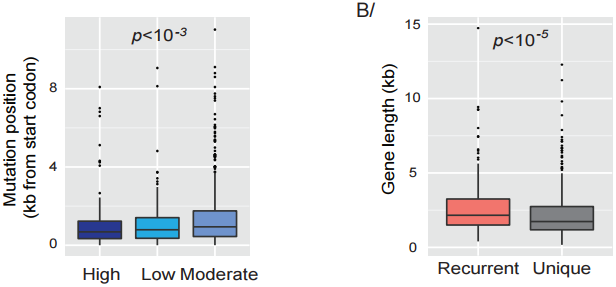
Distribution of high impact mutations. **A-** Enrichment of disruptive mutations (high impact) near the beginning of the gene. The significance of the difference between the three boxplots representing the distribution of the mutations within genes was estimated using a Wilcoxon rank-sum test. **B-** Distribution of the gene size between genes found recurrently mutated and genes found with only one mutation. The significance of the difference between the two boxplots representing the distribution of the sizes of genes in the two sets was estimated using a Wilcoxon rank-sum test.

**Figure S4.**
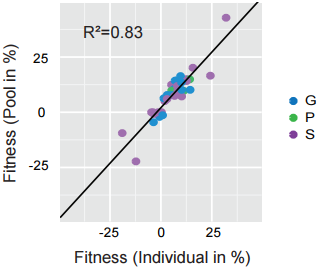
Fitnesses of 51 mutant strains measured in pool by barseq and in pairwise competitive assays. Because the fitness measured in the pooled experiment corresponds to the fitness relative to the population's mean fitness, we compared the pooled fitness data of 51 strains to individual fitness assays and found a strong positive correlation. Pearson's correlation coefficient R^2^= 0.83. G: Glucose-limited, S: Sulfate-limited and P: Phosphate-limited conditions.

**Figure S5.**
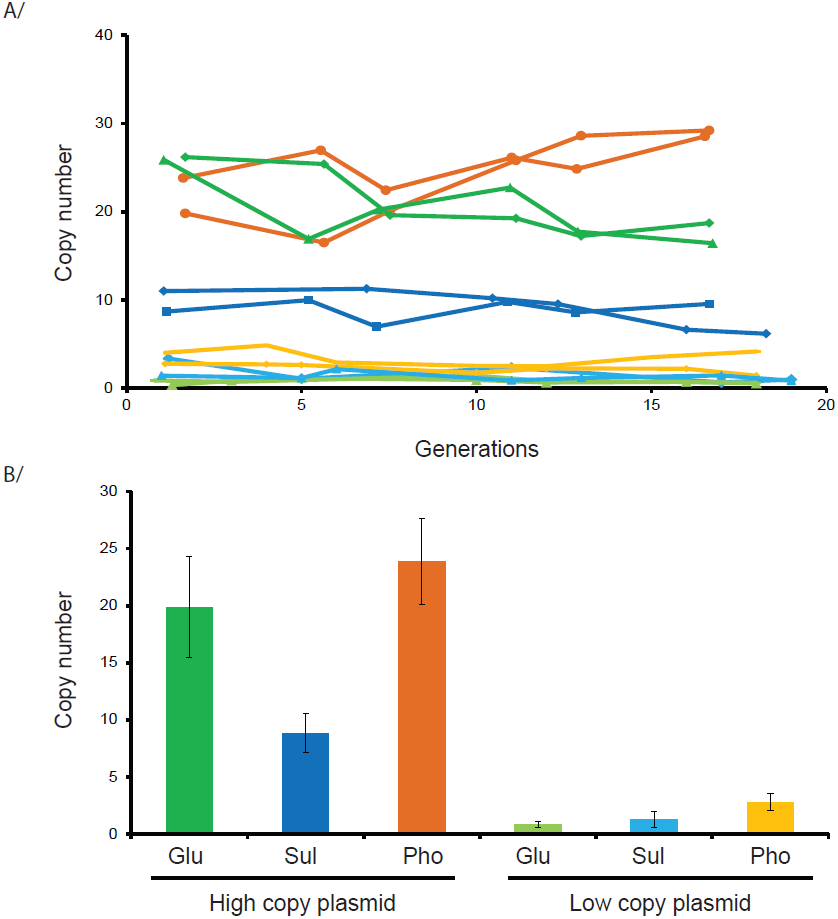
Fluctuations of the copy number of the plasmids monitored by qPCR on population samples over time. **A-**. Each color corresponds to a condition as described in panel B. **B-** Average of the plasmid copy number for both the high copy and the low copy plasmid collections grown for 20 generations in glucose-limited, sulfate-limited and phosphate-limited conditions.

Table S1: Strains and strain collections used in this study.

Table S2: Fitness measurements from mutant collection competitions.

Table S3: Fitness measurements from barcoder collection competitions.

Table S4: Identities, frequencies, and predicted effects of mutations discovered in experimental evolution studies.

Table S5: Beneficial mutations from the mutant collection competitions.

Table S6: Beneficial mutations in evolved samples.

Table S7: Fitness measurements from individual competition experiments vs pooled experiments.

Table S8: Primers used in this study.

Table S9: Barcode sequences used in this study.

Table S10: Summary statistics for barcode sequencing experiments.

Supplementary File 1: Perl script for demultiplexing sequencing files.

Supplementary File 2: C script used for barcode assignment.

Supplementary File 3: R script used for linear regression for fitness calculations.

## References

1. Alexandrov LB, Nik-Zainal S, Wedge DC, Aparicio SA, Behjati S, et al. (2013) Signatures of mutational processes in human cancer. Nature 500, 415–421.

2. Network TTCGAR (2013) Comprehensive genomic characterization defines human glioblastoma genes and core pathways. Nature 494, 506.

3. Forbes SA, Bhamra G, Bamford S, Dawson E, Kok C, et al. (2008) The Catalogue of Somatic Mutations in Cancer (COSMIC). Curr Protoc Hum Genet Chapter 10, Unit 10 11.

4. Cancer Genome Atlas N (2012) Comprehensive molecular characterization of human colon and rectal cancer. Nature 487, 330–337.

5. Lawrence MS, Stojanov P, Polak P, Kryukov GV, Cibulskis K, et al. (2013) Mutational heterogeneity in cancer and the search for new cancer-associated genes. Nature 499, 214–218.

6. Willingham AT, Deveraux QL, Hampton GM, Aza-Blanc P (2004) RNAi and HTS: exploring cancer by systematic loss-of-function. Oncogene 23, 8392–8400.

7. Willecke M, Toggweiler J, Basler K (2011) Loss of PI3K blocks cell-cycle progression in a *Drosophila* tumor model. Oncogene 30, 4067–4074.

8. Zender L, Xue W, Zuber J, Semighini CP, Krasnitz A, et al. (2008) An oncogenomics-based in vivo RNAi screen identifies tumor suppressors in liver cancer. Cell 135, 852–864.

9. Turner TL, Stewart AD, Fields AT, Rice WR, Tarone AM (2011) Population-based resequencing of experimentally evolved populations reveals the genetic basis of body size variation in *Drosophila melanogaster*. PLoS Genet 7, e1001336.

10. Lang GI, Rice DP, Hickman MJ, Sodergren E, Weinstock GM, et al. (2013) Pervasive genetic hitchhiking and clonal interference in forty evolving yeast populations. Nature Aug 29;500(7464):571–4.: 571–574.

11. Kvitek DJ, Sherlock G (2013) Whole genome, whole population sequencing reveals that loss of signaling networks is the major adaptive strategy in a constant environment. PLoS Genet 9, e1003972.

12. Hong J, Gresham D (2014) Molecular specificity, convergence and constraint shape adaptive evolution in nutrient-poor environments. PLoS Genet 10, e1004041.

13. Zhu YO, Siegal ML, Hall DW, Petrov DA (2014) Precise estimates of mutation rate and spectrum in yeast. Proc Natl Acad Sci U S A 2014 Jun 3;111(22): E2310–2318.

14. Barrick JE, Yu DS, Yoon SH, Jeong H, Oh TK, et al. (2009) Genome evolution and adaptation in a long-term experiment with *Escherichia coli*. Nature 461, 1243–1247.

15. Lee MC, Marx CJ (2013) Synchronous waves of failed soft sweeps in the laboratory: remarkably rampant clonal interference of alleles at a single locus. Genetics 193, 943–952.

16. Kao KC, Sherlock G (2008) Molecular characterization of clonal interference during adaptive evolution in asexual populations of *Saccharomyces cerevisiae*. Nat Genet 40, 1499–1504.

17. Tenaillon O, Rodriguez-Verdugo A, Gaut RL, McDonald P, Bennett AF, et al. (2012) The molecular diversity of adaptive convergence. Science 335, 457–461.

18. Herron MD, Doebeli M (2013) Parallel evolutionary dynamics of adaptive diversification in *Escherichia coli*. PLoS Biol 11, e1001490.

19. Dunham MJ, Badrane H, Ferea T, Adams J, Brown PO, et al. (2002) Characteristic genome rearrangements in experimental evolution of *Saccharomyces cerevisiae*. Proc Natl Acad Sci U S A 99, 16144–16149.

20. Blount ZD, Barrick JE, Davidson CJ, Lenski RE (2012) Genomic analysis of a key innovation in an experimental *Escherichia coli* population. Nature 489, 513–518.

21. Sniegowski PD, Gerrish PJ, Lenski RE (1997) Evolution of high mutation rates in experimental populations of *E. coli*. Nature 387, 703–705.

22. Gerstein AC, Lo DS, Otto SP (2012) Parallel genetic changes and nonparallel gene environment interactions characterize the evolution of drug resistance in yeast. Genetics 192, 241–252.

23. Chou HH, Berthet J, Marx CJ (2009) Fast growth increases the selective advantage of a mutation arising recurrently during evolution under metal limitation. PLoS Genet 5, e1000652.

24. Kvitek DJ, Sherlock G (2011) Reciprocal sign epistasis between frequently experimentally evolved adaptive mutations causes a rugged fitness landscape. PLoS Genet 7, e1002056.

25. Smith AM, Heisler LE, Mellor J, Kaper F, Thompson MJ, et al. (2009) Quantitative phenotyping via deep barcode sequencing. Genome Res 19, 1836–1842.

26. Delneri D, Hoyle DC, Gkargkas K, Cross EJ, Rash B, et al. (2008) Identification and characterization of high-flux-control genes of yeast through competition analyses in continuous cultures. Nat Genet 40, 113–117.

27. Sopko R, Huang D, Preston N, Chua G, Papp B, et al. (2006) Mapping pathways and phenotypes by systematic gene overexpression. Mol Cell 21, 319–330.

28. Makanae K, Kintaka R, Makino T, Kitano H, Moriya H (2013) Identification of dosage sensitive genes in *Saccharomyces cerevisiae* using the genetic tug-of-war method. Genome Res 23, 300–311.

29. Gelperin DM, White MA, Wilkinson ML, Kon Y, Kung LA, et al. (2005) Biochemical and genetic analysis of the yeast proteome with a movable ORF collection. Genes Dev 19, 2816–2826.

30. Costanzo M, Baryshnikova A, Bellay J, Kim Y, Spear ED, et al. (2010) The genetic landscape of a cell. Science 327, 425–431.

31. Douglas AC, Smith AM, Sharifpoor S, Yan Z, Durbic T, et al. (2012) Functional analysis with a barcoder yeast gene overexpression system. G3 (Bethesda) 2, 1279–1289.

32. Ho CH, Magtanong L, Barker SL, Gresham D, Nishimura S, et al. (2009) A molecular barcoded yeast ORF library enables mode-of-action analysis of bioactive compounds. Nat Biotechnol 27, 369–377.

33. Magtanong L, Ho CH, Barker SL, Jiao W, Baryshnikova A, et al. (2011) Dosage suppression genetic interaction networks enhance functional wiring diagrams of the cell. Nat Biotechnol 29, 505–511.

34. Tong AH, Boone C (2006) Synthetic genetic array analysis in *Saccharomyces cerevisiae*. Methods Mol Biol 313, 171–192.

35. Yan Z, Costanzo M, Heisler LE, Paw J, Kaper F, et al. (2008) Yeast Barcoders: a chemogenomic application of a universal donor-strain collection carrying bar-code identifiers. Nat Methods 5, 719–725.

36. Hillenmeyer ME, Ericson E, Davis RW, Nislow C, Koller D, et al. (2010) Systematic analysis of genome-wide fitness data in yeast reveals novel gene function and drug action. Genome Biol 11, R30.

37. Hillenmeyer ME, Fung E, Wildenhain J, Pierce SE, Hoon S, et al. (2008) The chemical genomic portrait of yeast: uncovering a phenotype for all genes. Science 320, 362–365.

38. Suzuki Y, St Onge RP, Mani R, King OD, Heilbut A, et al. (2011) Knocking out multigene redundancies via cycles of sexual assortment and fluorescence selection. Nat Methods 8, 159–164.

39. Qian W, Ma D, Xiao C, Wang Z, Zhang J (2012) The genomic landscape and evolutionary resolution of antagonistic pleiotropy in yeast. Cell Rep 2, 1399–1410.

40. Paquin C, Adams J (1983) Frequency of fixation of adaptive mutations is higher in evolving diploid than haploid yeast populations. Nature 302, 495–500.

41. Otto S (1994) The role of deleterious and beneficial mutations in the evolution of ploidy levels. Lectures on Mathematics in the Life Sciences 25.

42. Gresham D, Desai MM, Tucker CM, Jenq HT, Pai DA, et al. (2008) The repertoire and dynamics of evolutionary adaptations to controlled nutrient-limited environments in yeast. PLoS Genet 4, e1000303.

43. Payen C, Di Rienzi SC, Ong GT, Pogachar JL, Sanchez JC, et al. (2014) The Dynamics of Diverse Segmental Amplifications in Populations of *Saccharomyces cerevisiae* Adapting to Strong Selection. G3 (Bethesda) 4, 399–409.

44. Culotta VC, Lin SJ, Schmidt P, Klomp LW, Casareno RL, et al. (1999) Intracellular pathways of copper trafficking in yeast and humans. Adv Exp Med Biol 448, 247–254.

45. Portnoy ME, Liu XF, Culotta VC (2000) *Saccharomyces cerevisiae* expresses three functionally distinct homologues of the nramp family of metal transporters. Mol Cell Biol 20, 7893–7902.

46. Wenger JW, Piotrowski J, Nagarajan S, Chiotti K, Sherlock G, et al. (2011) Hunger artists: yeast adapted to carbon limitation show trade-offs under carbon sufficiency. PLoS Genet 7, e1002202.

47. Wray GA (2007) The evolutionary significance of cis-regulatory mutations. Nat Rev Genet 8, 206–216.

48. Cingolani P, Platts A, Wang le L, Coon M, Nguyen T, et al. (2012) A program for annotating and predicting the effects of single nucleotide polymorphisms, SnpEff: SNPs in the genome of *Drosophila melanogaster* strain w1118; iso-2; iso-3. Fly (Austin) 6, 80–92.

49. Gerstein AC, Kuzmin A, Otto SP (2014) Loss-of-heterozygosity facilitates passage through Haldane's sieve for *Saccharomyces cerevisiae* undergoing adaptation. Nat Commun 5, 3819.

50. Mwenifumbo JC, Marra MA (2013) Cancer genome-sequencing study design. Nat Rev Genet 14, 321–332.

51. Gonzalez-Perez A, Lopez-Bigas N (2012) Functional impact bias reveals cancer drivers. Nucleic Acids Res 40, e169.

52. Behjati S, Tarpey PS, Sheldon H, Martincorena I, Van Loo P, et al. (2014) Recurrent PTPRB and PLCG1 mutations in angiosarcoma. Nat Genet 46, 376–379.

53. Notley-McRobb L, Ferenci T (2000) Experimental analysis of molecular events during mutational periodic selections in bacterial evolution. Genetics 156, 1493–1501.

54. Poelwijk FJ, Kiviet DJ, Weinreich DM, Tans SJ (2007) Empirical fitness landscapes reveal accessible evolutionary paths. Nature 445, 383–386.

55. O'Connell KF, Baker RE (1992) Possible cross-regulation of phosphate and sulfate metabolism in *Saccharomyces cerevisiae*. Genetics 132, 63–73.

56. Ben-Shitrit T, Yosef N, Shemesh K, Sharan R, Ruppin E, et al. (2012) Systematic identification of gene annotation errors in the widely used yeast mutation collections. Nat Methods 9, 373–378.

57. Yaniv M (2014) Chromatin remodeling: from transcription to cancer. Cancer Genet.

58. Chung N, Jenkins G, Hannun YA, Heitman J, Obeid LM (2000) Sphingolipids signal heat stress-induced ubiquitin-dependent proteolysis. J Biol Chem 275, 17229–17232.

59. Pavlov YI, Shcherbakova PV (2010) DNA polymerases at the eukaryotic fork-20 years later. Mutat Res 685, 45–53.

60. Lynch M (2008) The cellular, developmental and population-genetic determinants of mutation-rate evolution. Genetics 180, 933–943.

61. Gerstein AC (2013) Mutational effects depend on ploidy level: all else is not equal. Biol Lett 9, 20120614.

62. Tang YC, Amon A (2013) Gene copy-number alterations: a cost-benefit analysis. Cell 152, 394–405.

63. Torres EM, Sokolsky T, Tucker CM, Chan LY, Boselli M, et al. (2007) Effects of aneuploidy on cellular physiology and cell division in haploid yeast. Science 317, 916–924.

64. de Visser JA, Krug J (2014) Empirical fitness landscapes and the predictability of evolution. Nat Rev Genet 15, 480–490.

65. Hall DW, Mahmoudizad R, Hurd AW, Joseph SB (2008) Spontaneous mutations in diploid Saccharomyces cerevisiae: another thousand cell generations. Genet Res (Camb) 90, 229–241.

66. Fowler DM, Araya CL, Fleishman SJ, Kellogg EH, Stephany JJ, et al. (2010) High resolution mapping of protein sequence-function relationships. Nat Methods 7, 741–746.

67. Thompson O, Edgley M, Strasbourger P, Flibotte S, Ewing B, et al. (2013) The million mutation project: a new approach to genetics in *Caenorhabditis elegans*. Genome Res 23, 1749–1762.

68. Bloom-Ackermann Z, Navon S, Gingold H, Towers R, Pilpel Y, et al. (2014) A comprehensive tRNA deletion library unravels the genetic architecture of the tRNA pool. PLoS Genet 10, e1004084.

69. Pepin KM, Wichman HA (2007) Variable epistatic effects between mutations at host recognition sites in phiX174 bacteriophage. Evolution 61, 1710–1724.

70. Elena SF, Lenski RE (1997) Test of synergistic interactions among deleterious mutations in bacteria. Nature 390, 395–398.

71. Jasnos L, Korona R (2007) Epistatic buffering of fitness loss in yeast double deletion strains. Nat Genet 39, 550–554.

72. Kryazhimskiy S, Rice DP, Jerison ER, Desai MM (2014) Microbial evolution. Global epistasis makes adaptation predictable despite sequence-level stochasticity. Science 344, 1519–1522.

73. Costanzo M, Baryshnikova A, Bellay J, Kim Y, Spear ED, et al. The genetic landscape of a cell. Science 327, 425–431.

74. Tong AH, Lesage G, Bader GD, Ding H, Xu H, et al. (2004) Global mapping of the yeast genetic interaction network. Science 303, 808–813.

75. Supek F, Minana B, Valcarcel J, Gabaldon T, Lehner B (2014) Synonymous mutations frequently act as driver mutations in human cancers. Cell 156, 1324–1335.

76. Hoekstra HE, Coyne JA (2007) The locus of evolution: evo devo and the genetics of adaptation. Evolution 61, 995–1016.

77. Gresham D, Usaite R, Germann SM, Lisby M, Botstein D, et al. (2010) Adaptation to diverse nitrogen-limited environments by deletion or extrachromosomal element formation of the *GAP1* locus. Proc Natl Acad Sci U S A 107, 18551–18556.

78. Hoffman CS, Winston F (1987) A ten-minute DNA preparation from yeast efficiently releases autonomous plasmids for transformation of *Escherichia coli*. Gene 57, 267–272.

